# TDP-43 Aggregate Seeding Impairs Autoregulation and Causes TDP-43 Dysfunction

**DOI:** 10.1101/2025.02.11.637743

**Authors:** Lohany Dias Mamede, Miwei Hu, Amanda R. Titus, Jaime Vaquer-Alicea, Rachel L. French, Marc I. Diamond, Timothy M. Miller, Yuna M. Ayala

## Abstract

The aggregation, cellular mislocalization and dysfunction of TDP-43 are hallmarks of multiple neurodegenerative disorders. We find that inducing TDP-43 aggregation through prion-like seeding gradually diminishes normal TDP-43 nuclear localization and function. Aggregate-affected cells show signature features of TDP-43 loss of function, such as DNA damage and dysregulated TDP-43-target expression. We also observe strong activation of TDP-43-controlled cryptic exons in cells, including human neurons treated with proteopathic seeds. Furthermore, aggregate seeding impairs TDP-43 autoregulation, an essential mechanism controlling TDP-43 homeostasis. Interestingly, proteins that normally interact with TDP-43 are not recruited to aggregates, while other factors linked to TDP-43 pathology, including Ataxin 2, specifically colocalize to inclusions and modify seeding-induced aggregation. Our findings indicate that TDP-43 aggregation, mislocalization and loss of function are strongly linked and suggest that disruption of TDP-43 autoregulation establishes a toxic feed-forward mechanism that amplifies aggregation and may be central in mediating this pathological connection.

## INTRODUCTION

The Transactive Response DNA binding protein (TDP-43) is an essential RNA binding protein that regulates gene expression through RNA processing and is strongly linked to multiple neurodegenerative disorders, collectively referred to as TDP-43 proteinopathies. TDP-43 pathology is characterized by protein aggregation, most often in the cytoplasm of neurons and glial cells, and depletion from its normal nuclear distribution^1^. TDP-43 is an RNA binding protein most well-characterized for regulating alternative splicing and alternative polyadenylation^2, 3, 4, 5, 6^. Recent findings have underscored the involvement of loss of TDP-43 function in disease, as dysregulation of TDP-43 target genes is observed in patients affected by TDP-43 proteinopathies^7, 8, 9, 10, 11, 12^. Collectively, these observations strongly suggest that TDP-43 misfolding and aggregation are correlated with the loss of protein function. However, the mechanisms connecting these two major components of TDP-43 pathology remain unclear.

TDP-43 pathology is hallmark feature of neurotoxicity and neurodegeneration in almost all amyotrophic lateral sclerosis (ALS) and ∼50% of frontotemporal lobar degeneration (FTLD-TDP) and the ensuing frontotemporal dementia (FTD)^1, 13^. TDP-43 inclusions also characterize an Alzheimer’s disease (AD)-associated disorder, limbic-predominant age-related TDP-43 encephalopathy (LATE), that affects nearly a quarter of individuals over 85 years of age^14^. In addition, TDP-43 is a comorbid feature of greater than 50% of AD cases, correlated with more aggressive memory loss and brain atrophy^15, 16, 17, 18^. Similarly, TDP-43 aggregates are major components upon traumatic brain injury and chronic traumatic encephalopathy^19, 20^. A common feature among TDP-43-linked pathologies and other neurodegenerative proteinopathies is the propagation of protein toxicity through prion-like aggregate seeding. This process is mediated by a self-perpetuating mechanism, where misfolded protein induces the aggregation of its soluble form. Clinicopathological data shows the progressive spread of TDP-43 pathology in patient brain and spinal cord^14, 17, 21, 22^, supporting cell-to-cell transmission of TDP-43 aggregates. TDP-43 aggregate seeding has been demonstrated experimentally by showing that aggregates generated from purified TDP-43 are seeding competent in cell culture^23, 24^. Additionally, insoluble extracts rich in phosphorylated TDP-43 (pTDP-43) initiate de novo aggregation in the brain of a TDP-43 mouse model^25^. Aggregate seeding efficiency is enhanced by exposing the fibrillar C-terminal domain (CTD) following limited proteolysis of TDP-43 aggregates^26^. This is a mostly disordered domain referred to as the prion-like domain for its sequence similarity to yeast prion proteins^27^. The CTD forms the core filament structure derived from FTD patient brain, specifically between amino acids 282-360^28^ (**Fig. 1A**). Collectively, these observations underscore the critical role of this region during misfolding and transmission of disease-relevant aggregates. The mechanisms of proteopathic seed uptake, propagation and the consequences of aggregate seeding on TDP-43 cellular function remain poorly understood. Yet, elucidation of these processes is critical to define mechanisms underlying disease and to identify potential therapeutic strategies.

**Figure 1.**
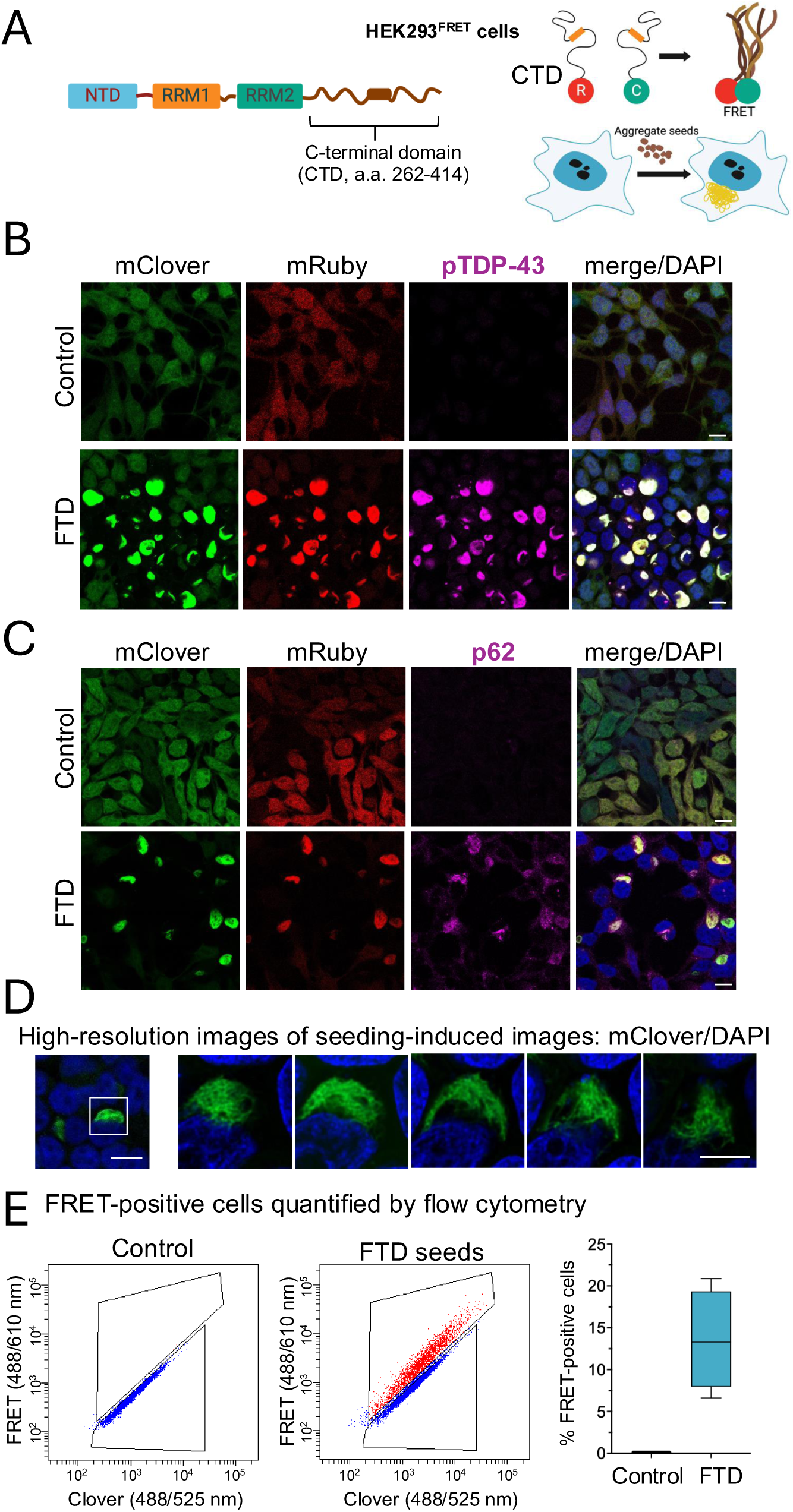
Disease-derived extract strongly activates TDP-43 aggregate seeding in a sensitive FRET-based biosensor. A) HEK293 cells stably expressing the C-terminal prion like, disordered domain of TDP-43 (CTD) fused to mClover3 and mRuby3 FRET pairs (HEK293^FRET^) used as a reporter of TDP-43 aggregation upon seeding with FTD extract prepared as in Fig. S1. B) Visualization of de novo CTD aggregation in HEK293^FRET^ treated with FTD seeds or neurological unaffected tissue extract, used as control. Antibodies for phosphorylated TDP-43 (pTDP-43) and p62, shown in (C), were used as markers of pathological TDP-43 aggregates. Images were obtained by confocal microscopy. DAPI staining is shown to highlight nuclei in the overlay image. Scale, 10 μm. D) High resolution images of FTD-treaded cell aggregate showing z-stacks with DAPI overlay generated with stimulated emission depletion (STED) microscopy. E) Flow cytometry to quantify FRET-positive and FRET-negative HEK293^FRET^ cells six days after treatment with FTD seeds and control extract. The proportion of FRET-positive cells in each group was quantified as percent FRET-positive cells compared to total, n>3.

Experimental methods to recapitulate TDP-43 aggregation and loss of function in the same model have been challenging to develop and remain rare. To induce TDP-43 aggregation, common models rely on overexpression of wild-type or mutant TDP-43 or chemical treatment to induce oxidative stress or other proteotoxic conditions. These models trigger generalized disruption of cell function and undermine insight into disease-relevant mechanisms. Furthermore, these experimental conditions rarely lead to TDP-43 depletion, and their effect on TDP-43 function has not been widely reported. Other assays to investigate TDP-43 dysfunction rely on knocking-out/down TDP-43 expression, however, these conditions lack the contribution of TDP-43 mislocalization and aggregation. Thus, to establish cellular models combining both TDP-43 aggregation and loss of protein function mechanisms that overcome the limitations of previous methods, we explored systems based on TDP-43 prion-like aggregate seeding. Using FTD brain-derived seeds, we find that aggregate seeding greatly impacts TDP-43 localization and leads to gradual loss of function. Importantly, aggregate seeding disrupts TDP-43 autoregulation, increasing TDP-43 expression. Our model recapitulates key features of TDP-43 pathology and provides evidence that aggregation, mislocalization and loss of function mechanisms are interconnected.

## RESULTS

### TDP-43 aggregate seeding triggers de novo aggregation and gradual TDP-43 mislocalization

To quantify, isolate and characterize cells affected by TDP-43 aggregates induced by intracellular seeding, we employed a human embryonic kidney cell line (HEK293) that reports TDP-43 aggregation based on Förster resonance energy transfer (FRET)^29^. This cell line stably expresses the TDP-43 C-terminal domain (CTD, amino acids 263-414) fused N-terminally to FRET pairs mClover3 and mRuby3 (HEK293^FRET^) (**Fig. 1A**). To investigate TDP-43 aggregate seeding in the context of disease-derived proteopathic seeds, we isolated the insoluble fraction of autopsy brain from FTLD-TDP patients subtypes A and B^30, 31, 32, 33, 34, 35, 36, 37^. We prepared sarkosyl-insoluble fractions upon sequential extraction based on previously established methods^38^. Herein, we refer to this extract as FTD seeds. These samples were enriched for total TDP-43 and a marker of pathological inclusions, phosphorylated TDP-43 (pTDP-43, Ser409/410)^39^ (**Fig. S1**). As control, we processed brain tissue from neurologically unaffected patient, which did not show significant accumulation of pTDP-43. Treatment of HEK293^FRET^ cells with FTD seeds induced the formation of cytoplasmic inclusions composed of mClover and mRuby-CTDs that were detected by antibodies specific for pTDP-43 (**Fig. 1B**) and the ubiquitin binding protein 62/sequestosome 1 (p62/SQTM1) (**Fig. 1C**), which is often associated with TDP-43 inclusions in disease^30, 40^. The seeding-induced aggregates often showed filamentous morphology, as seen by high-resolution microscopy (**Fig. 1D**). Analogous cell lines expressing only mClover- or mRuby-fused CTD showed similar aggregate seeding behavior (**Fig. S2**). We quantified the number of cells that developed aggregates upon seeding by flow cytometry and determined the ratio of FRET-positive cells relative to the total number of cells. Treatment with FTD seeds resulted in an average of 13% FRET-positive cells, compared to <1% for control tissue extract (**Fig. 1E**). These findings are consistent with the previously observed sensitivity of this reporter cell line to seeding with pre-formed aggregates of purified TDP-43 and aggregates derived from a mouse model of TDP-43 pathology in skeletal muscle^29, 41^. The significant increase in FRET activity upon treatment with FTD seeds, compared to control-treated cells in which this signal is nearly absent, suggests that homotypic reversible CTD interactions, such as phase separation^42, 43^, do not significantly contribute to the FRET signal detected in our analyses. Alternatively, the concentration of the CTD fragments in HEK293^FRET^ cells is not sufficient to generate these assemblies under the conditions tested. Intriguingly, while FTD seeds triggered aggregates with characteristic fibrillar structure, pre-formed aggregates from purified TDP-43 resulted in smaller and more compact de novo inclusions (**Fig. 1D, S3**).

We next investigated changes in the localization of endogenous TDP-43 following seeding-induced aggregation. To specifically detect endogenous TDP-43, we used an antibody recognizing an N-terminal domain epitope absent in mClover- and mRuby-CTD constructs. Despite the accumulation of cytoplasmic aggregates three days post-treatment with FTD seeds, endogenous TDP-43 localization remained largely unaffected (**Fig. 2A**, arrows). In contrast, by six days post-seeding, we observed colocalization of endogenous TDP-43 with CTD cytoplasmic inclusions, accompanied by marked reduction of its nuclear localization (**Fig. 2B**). To quantify depletion of nuclear TDP-43 distribution in cells treated with FTD seeds, we measured the mean fluorescence intensity of endogenous TDP-43 in the region of interest defined as the nucleus, based on DAPI staining. From this analysis we found that endogenous TDP-43 nuclear levels decreased significantly by approximately 86% percent in cells affected by cytoplasmic aggregates, compared to cells lacking cytoplasmic aggregates (**Fig. 2C**). These findings indicate that cytoplasmic CTD aggregation initiated by seeding sequesters endogenous TDP-43, leading to its gradual depletion from the nucleus, and recapitulates key features of neuronal TDP-43 pathology in patients^1^.

**Figure 2:**
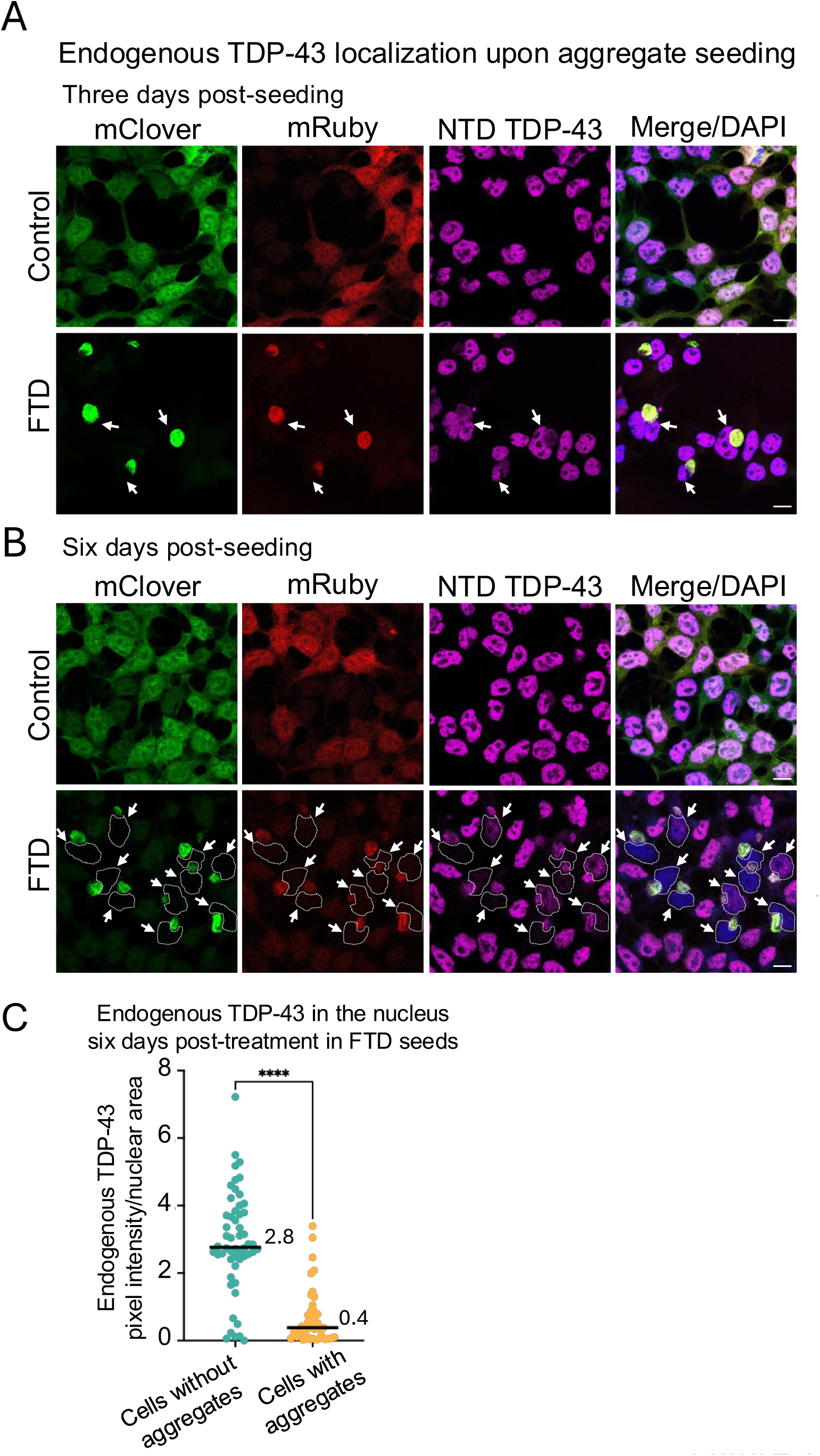
Aggregate seeding induces co-aggregation and gradual depletion of nuclear endogenous TDP-43. A) Immunofluorescence of HEK293^FRET^ cells treated with control and FTD-derived extract, probed with a TDP-43 antibody targeting an N-terminal domain epitope to detect endogenous protein only. Cells were imaged three and six days post-treatment in (B), using confocal microscopy. Arrows indicate cytoplasmic aggregates. DAPI staining highlights nuclei in the overlay (merge) image. The nuclei of aggregate-affected cells treated with FTD seeds are outlined with white dashed lines. C) Fluorescence intensity of endogenous TDP-43 was measured in the nuclear compartment of cells six days post-treatment with FTD seeds, defined by DAPI staining. ImageJ was used to quantify pixel intensity per area of >200 cells from n>3. Calculated median values are shown, ****p<0.0001, Mann-Whitney test.

### Aggregate seeding results in loss of TDP-function in affected cells

Upon observing the loss of nuclear TDP-43 in aggregate-affected cells (**Fig. 2B, C**), we examined whether this correlates with loss of TDP-43 function. First, we compared the impact on genomic stability, as TDP-43 downregulation increases accumulation of DNA breaks in human cells and neurons, according to well-established data^44, 45, 46^. Six days post-seeding, phosphorylated γH2AX, a marker of DNA breaks, increased and was detected in 50 ± 7% of TDP-43 aggregate-positive cells (>15 γH2AX foci per nucleus) (**Fig. 3A**). In contrast, control-treated cells showed 4% γH2AX-positive nuclei. These observations were consistent with reduced nuclear TDP-43 levels and indicated impaired protein function cells accumulating aggregates. To further examine TDP-43 activity in these cells, we measured mRNA transcript expression and processing of TDP-43-regulated targets. Based on previous studies in human cells, we selected transcripts controlled by TDP-43 through different posttranscriptional processing mechanisms. TDP-43 binds to the 3’UTR of *SMCA1* and *GXYLT1* controlling the usage of alternative polyadenylation poly(A) sites, inhibiting gene expression^6^ (**Fig. 3B**). We sorted FRET-positive cells to separate them from FRET-negative cells via FACS at six days post-seeding with FTD seeds (**Fig. 1D**). Using quantitative PCR, we measured mRNA expression in cells treated with FTD seeds, control-derived extracts and non-treated control. FRET-positive cells showed approximately 2-fold or greater significant increase in mRNA expression, compared to FRET-negative cells (**Fig. 3B**). Cells treated with control tissue extract or non-treated control were also sorted, resulting in mostly FRET-negative cells. Actin B mRNA (*ACTB*) was used as reference for a transcript not regulated by TDP-43 and this showed no significant differences in expression between groups. Similar analysis at earlier time points, three days post-seeding, when approximately 4% of cells treated with FTD seeds were FRET-positive, showed no significant changes in *SMCA1*, *GXYLT1* expression between FRET-positive and negative groups (not shown).

**Figure 3:**
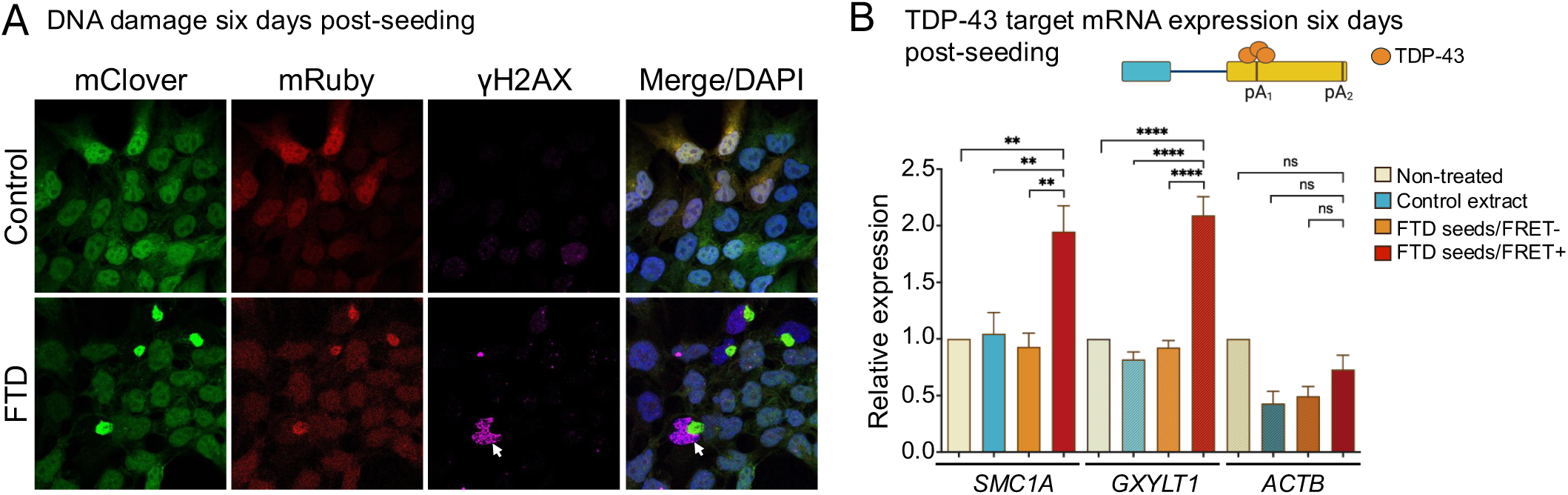
Seeding-induced aggregation causes loss of TDP-43 function. A) Immunofluorescence obtained by confocal microscopy to detect phosphorylated γH2AX, a marker of DNA breaks, in cells six days post-treatment with control of FTD-seeds. Arrow points to the accumulation of phosphorylated γH2AX in aggregate-affected cells. Scale, 10 μm. B) TDP-43 regulates 3’ UTR processing and polyadenylation choice via direct recruitment to transcripts, as illustrated, where TDP-43 binding inhibits mRNA expression of *SMC1A and GXYLT1*. HEK293^FRET^ cells were collected six days post-treatment with FTD seeds, control extract and non-treated control, and sorted via FACS into FRET-positive (FRET+) and FRET-negative (FRET-) cells. Relative *SMC1A* and *GXYLT1* mRNA levels were measured by quantitative real-time PCR (qPCR). *ACTB* served as non-TDP-43-regulated transcript control. Graphs represent mean values, SEM of n>3. **p<0.005, ****p<0.0001, ns, non-significant, Mann-Whitney test.

In addition to regulating constitutive splicing of hundreds of genes, TDP-43 controls splicing of cryptic exons (CEs)–which constitute untranslated/intronic regions that are normally excluded from mature mRNA– by enhancing or inhibiting their inclusion^7^. Recent studies show the inclusion of TDP-43 target CEs in ALS-FTD cases and AD, consistent with loss of TDP-43 function^9, 10, 11, 12, 47, 48, 49, 50^. Thus, impaired regulation of CE splicing is viewed as a salient feature of TDP-43 dysfunction and disease, and abnormal mRNA products may be used as sensitive markers of TDP-43 pathology in patients. We investigated activation of CEs upon TDP-43 aggregate seeding in cases where TDP-43 inhibits CE inclusion, *HDGFL2* and *ARHGAP32* (**Fig. 4A**)^7, 51^. We observed CE activation in cells treated with FTD seeds starting at three days post-treatment, in which *ARHGAP32* CE splicing was significantly upregulated and *HDGFL2* showed an increasing trend. These non-sorted cells showed nearly two-fold greater CE inclusion compared to control. Strikingly, FRET-positive cells isolated six days post-seeding showed 150 and 1,000-fold greater *HDGFL2* and *ARHGAP32* CE inclusion, respectively, compared to FRET-negative and control-treated cells (**Fig. 4B**). This significant activation of CE inclusion starting at early timepoints after aggregate seeding, which dramatically increases in aggregate-affected cells, indicates that CE regulation of these targets, and perhaps others, are highly sensitive to decreasing TDP-43 levels.

**Figure 4:**
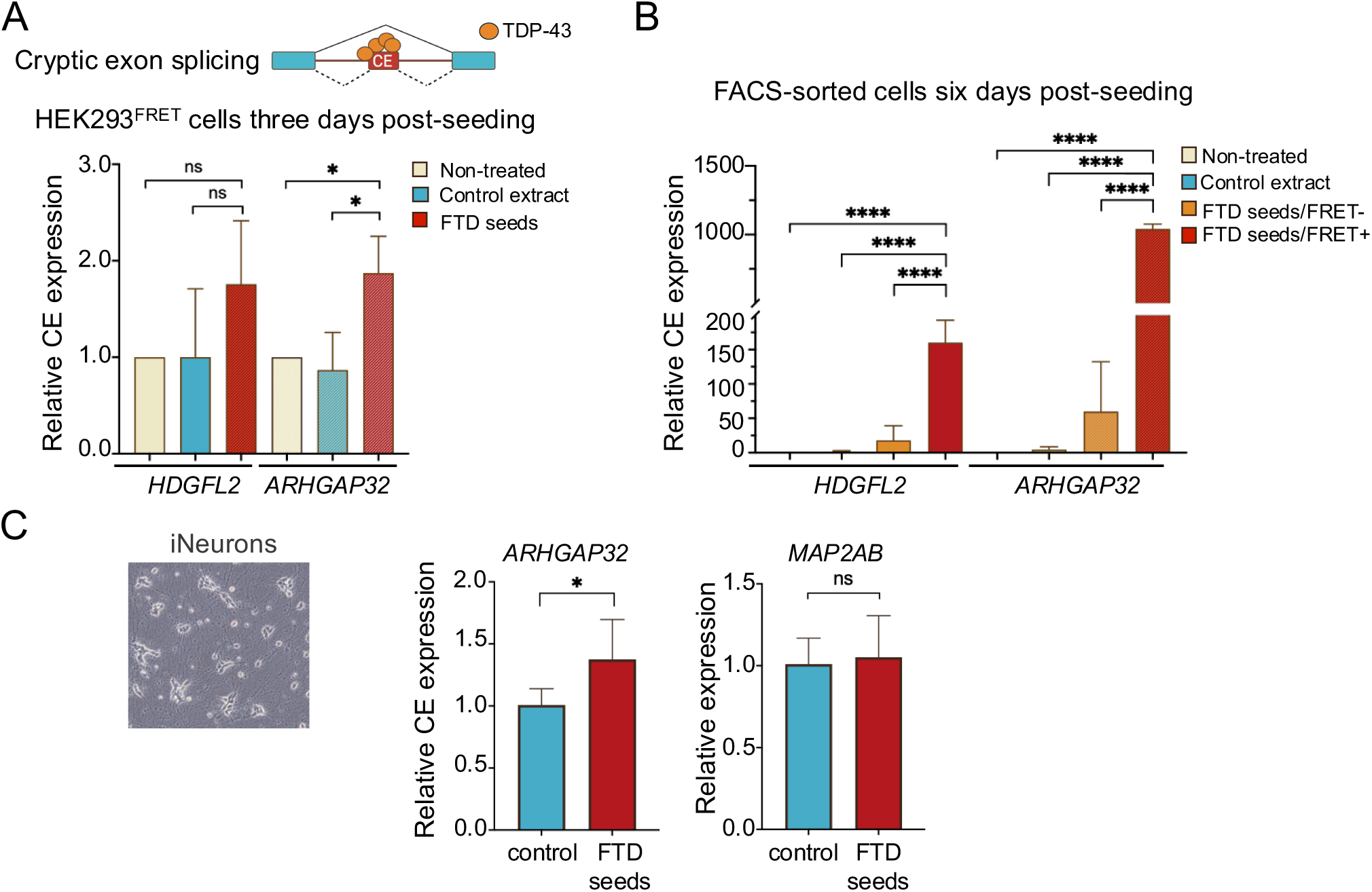
Aggregate seeding activates cryptic exon splicing of TDP-43-controlled targets. A) Cryptic exon (CE) splicing is normally suppressed by TDP-43 in *HDGFL2* and *ARHGAP32*. Relative CE inclusion was quantified by qPCR from HEK293^FRET^ cells collected three days post-treatment with FTD seeds, control-derived extract, or non-treated control. B) Relative *HDGFL2* and *ARHGAP32* CE inclusion quantified in cells sorted into FRET-positive (FRET+) and FRET-negative (FRET-) cells via FACS, six days post-seeding. C) Brightfield image of neurons derived from neurogenin-2 (Ngn2) inducible human induced pluripotent stem cells (iNeurons) 2-3 weeks after treatment with FTD seeds. Relative *ARHGAP32* CE inclusion was measured by qPCR. *MAP2AB* expression was measured to compare the abundance of microtubule-associated protein 2AB (MAP2AB)-positive neurons in FTD seed and control-treated samples. All graphs show mean values, SEM, n>3, *p<0.02, ****p<0.0001, ns, non-significant, Mann-Whitney test.

Next, we investigated whether TDP-43 aggregate seeding elicits loss of function in human neurons, as a more disease-relevant model. Induced pluripotent stem cells (iPSCs) with a doxycycline-inducible neurogenin-2 transgene integrated at the AAVS1 locus were differentiated into neurons (iNeurons)^52^ and treated with FTD seeds or control one week post-differentiation. iNeurons incubated for two to three weeks post-treatment with FTD seeds showed significant upregulation of *ARHGAP32* CE inclusion, relative to control treatment (**Fig. 4C**). Both treatment groups showed equivalent *MAP2AB* expression, a marker of neuronal differentiation and maturation. In all, our findings suggest that seeding-induced TDP-43 aggregation initiates a gradual loss of TDP-43 function, triggering defects in genomic stability and dysregulation of multiple targets that require protein activity.

### Aggregate seeding inhibits TDP-43 autoregulation impairing protein homeostasis

We next investigated whether TDP-43 functional defects triggered by aggregate seeding impact TDP-43 autoregulation, a critical process controlling TDP-43 homeostasis. TDP-43 regulates its own mRNA (*TARDBP*) and protein levels through a negative feedback loop that requires TDP-43 recruitment to the *TARDBP* 3’UTR within exon 6^5, 53^(**Fig. 5A**). This binding downregulates *TARDBP* mRNA expression by activating alternative poly(A) site usage, splicing and mRNA sequestration, which, combined, decrease protein expression^5, 53, 54^. Decreasing TDP-43 levels enhances usage of the proximal poly(A) site (pA_1_) and inhibits removal of alternative introns 6, 7 and 8 through splicing, overall increasing Short 3’UTR transcript expression and protein synthesis^54^. On the other hand, excess of TDP-43 protein levels increases the accumulation of multiple transcript isoforms that are targets of nonsense-mediated decay (NMD), as well as the Long 3’UTR which is a product of distal poly(A) site (pA_3_) selection and becomes sequestered in nuclear particles ^5, 54, 55, 56^. Maintaining physiological TDP-43 proteostasis through this evolutionarily conserved process is important for cell function as both diminished TDP-43 and elevated protein levels result in misregulated target expression. Moreover, abnormally increased TDP-43 concentration disrupts self-assembly and promotes aggregation (^57^for review). To determine whether aggregate seeding alters TDP-43 autoregulation, we compared *TARDBP* mRNA expression in FRET-positive and FRET-negative HEK293^FRET^ cells treated with FTD seeds. We also sorted cells treated with control tissue extract as well as non-treated control. We found significant, greater than two-fold upregulation of *TARDBP* mRNA expression in FRET-positive, compared to FRET-negative cells treated with FTD seeds (**Fig. 5B**). This significant increase was also observed relative to the control cells, which were almost all FRET-negative. Similar results were obtained using probes that are common to Short and Long 3’UTR mRNA isoforms, Ex 2-3, and a probe found in Short 3’UTR but absent in the NMD-sensitive isoforms, which captures the TDP-43 3’UTR binding region (TDPBR) upstream of pA_1_ (Ex 6). In contrast, a probe detecting the region closely upstream of the polyA_3_, specific for the Long 3’ UTR isoform, did not show significant differences in expression. These results suggest that aggregate seeding specifically upregulates pA_1_ usage and expression of the Short 3’UTR isoform, consistent with a loss of active TDP-43 levels. Moreover, this expression profile is consistent with that observed in FTD/ALS patient-derived cells devoid of nuclear TDP-43^8, 54^ and previous evidence showing specific upregulation of Short 3’UTR upon TDP-43 knockdown^54^. Based on these observations, we postulate that TDP-43 aggregate seeding blocks autoregulation and initiates a toxic feed-forward loop, amplifying aggregation due to uncontrolled protein synthesis and greater aggregate seeding.

**Figure 5:**
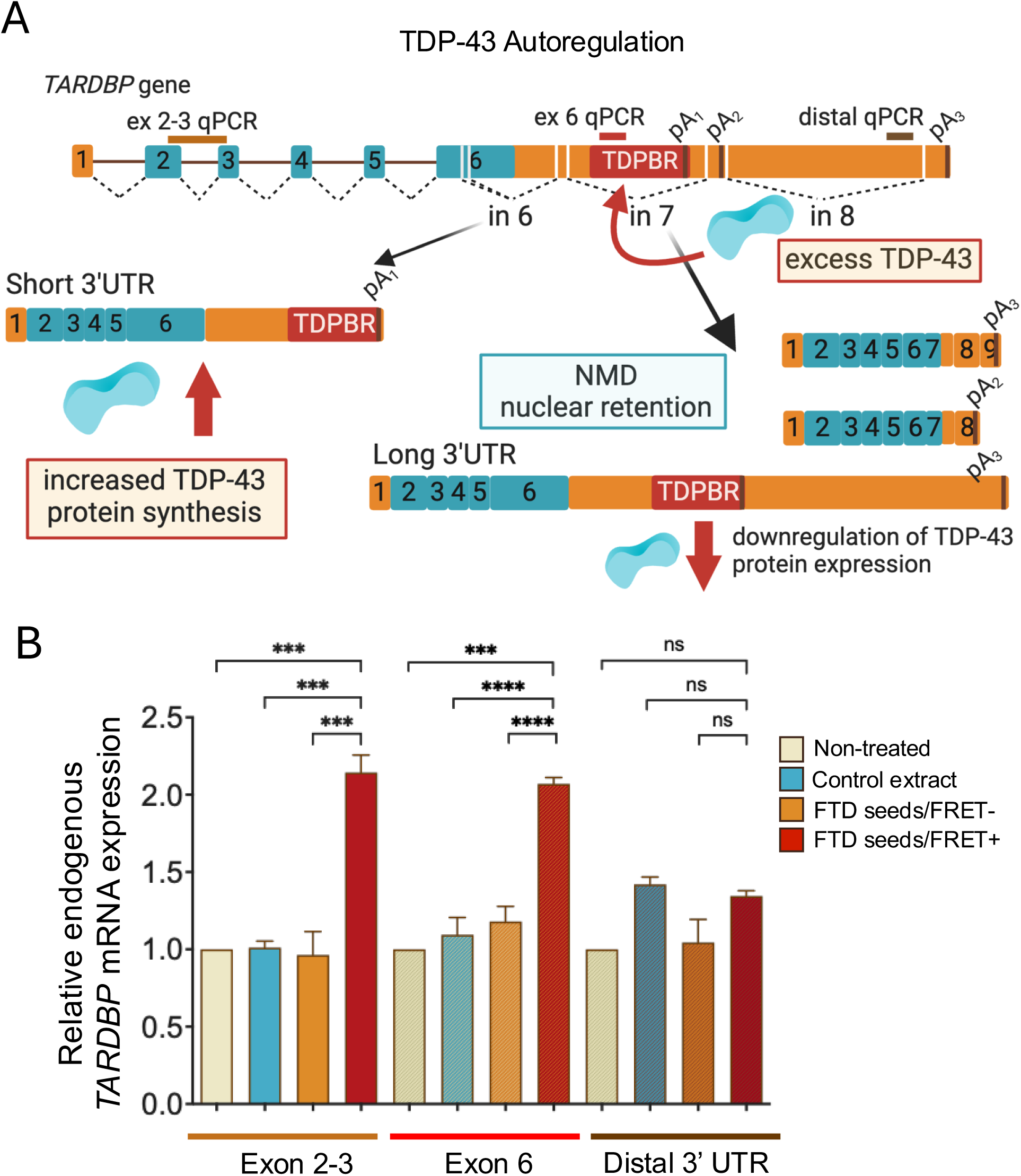
Aggregate seeding impairs TDP-43 autoregulation. A) Diagram illustrating mechanisms involved in TDP-43 autoregulation. TDP-43 protein binds to the TDP-43 Binding Region (TDPBR) within the 3’UTR of its own transcript, upstream of the proximal polyadenylation site (pA_1_) and near exon 6 splice sites (dashed lines indicate splicing events). TDP-43 recruitment promotes exon 6 splicing and distal polyadenylation site (pA_3_) usage. In the absence of TDP-43 binding, the Short 3’UTR isoform is upregulated, resulting in increased protein expression. qPCR probes are indicated. B) The relative expression of endogenous *TARDBP* mRNA was quantified by qPCR in FRET-positive (FRET+) and negative (FRET-) cells sorted via FACS, probing three regions of the transcript, as indicated. Mean values and SEM are shown for n>3, ***p<0.0002, ****p<0.0001, ns, non-significant, Mann Whitney test.

### TDP-43 aggregates specifically colocalize with modulators of aggregate seedin

Our findings showing strong co-localization of endogenous TDP-43 with cytoplasmic aggregates upon seeding (**Fig. 2B**) led us to explore whether this recruitment is specific to TDP-43 and whether these inclusions act as sinks for proteins that interact with TDP-43 under normal and pathological conditions. We focused on RNA binding proteins with disordered or prion-like domains that interact with TDP-43 during RNA processing, stress granule assembly, or disease-associated conditions: hnRNP A2/B1, hnRNP C1/C2, hnRNP H1, FUS, matrin 3, SFPQ, G3BP1, TIAR, HSPB1, and Ataxin 2^58, 59, 60, 61, 62^. Six days after treatment with FTD seeds we observed no significant colocalization of most proteins tested (**Figs. 6, S4A, B**). These findings suggest that seeding-induced aggregates in HEK293^FRET^ cells do not recruit proteins indiscriminately, even if these interact with TDP-43 under physiological conditions or during stress granule assembly. In contrast, ataxin 2 (*Atxn2*) and the small heat shock protein HSPB1 (*Hspb1*) strongly colocalized with de novo cytoplasmic aggregates (**Figs. 7A, S4C** arrows). Notably, *Atxn2* alters TDP-43 aggregation and toxicity in disease-relevant models^63, 64^ and upon induction of cytoplasmic TDP-43 misfolding in cells^65^. To test whether *Atxn2* expression influences seeding-induced TDP-43 aggregation, we reduced *Atxn2* expression by siRNA-mediated knockdown in HEK293^FRET^ cells, following treatment with FTD seeds. Decreasing *Atxn2* expression significantly reduced the accumulation of FRET-positive cells six days after treatment with FTD seeds (**Fig. 6B, S5**). We did not observe significant changes in the number of FRET-positive cells upon treatment with control, which accounted for <1% of all cells. These results suggest that *Atxn2* knockdown decreases seeding-induced TDP-43 aggregation, consistent with previous studies showing that *Atxn2* enhances TDP-43 aggregation and neurotoxicity in human cells and animal models^63, 66^. Overall, our findings suggest that TDP-43 prion-like aggregate seeding in our model triggers homotypic misfolding and co-aggregation with specific proteins. Moreover, the colocalization observed in our aggregation model may be linked to pathological interactions that impact disease progression.

**Figure 6.**
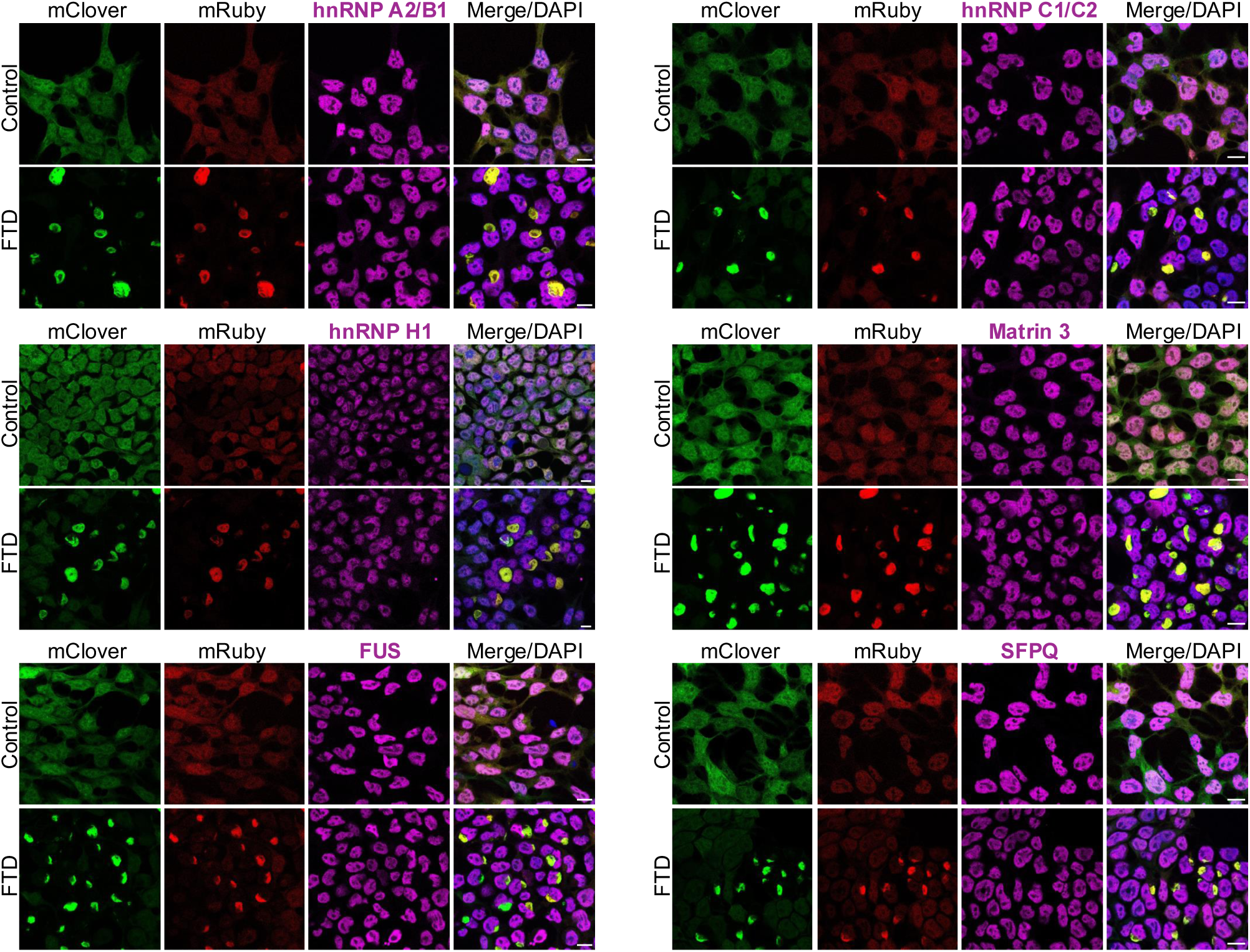
Protein partners that normally bind TDP-43 are not recruited to seeding-induced aggregates. Representative immunofluorescence images, obtained by confocal microscopy, of HEK293^FRET^ cells treated with FTD seeds or control extract. The antibodies probed RNA binding proteins that form complexes with TDP-43 under physiological conditions. Scale, 10 μm, n>3. DAPI staining was used to highlight nuclei in the overlay image.

**Figure 7.**
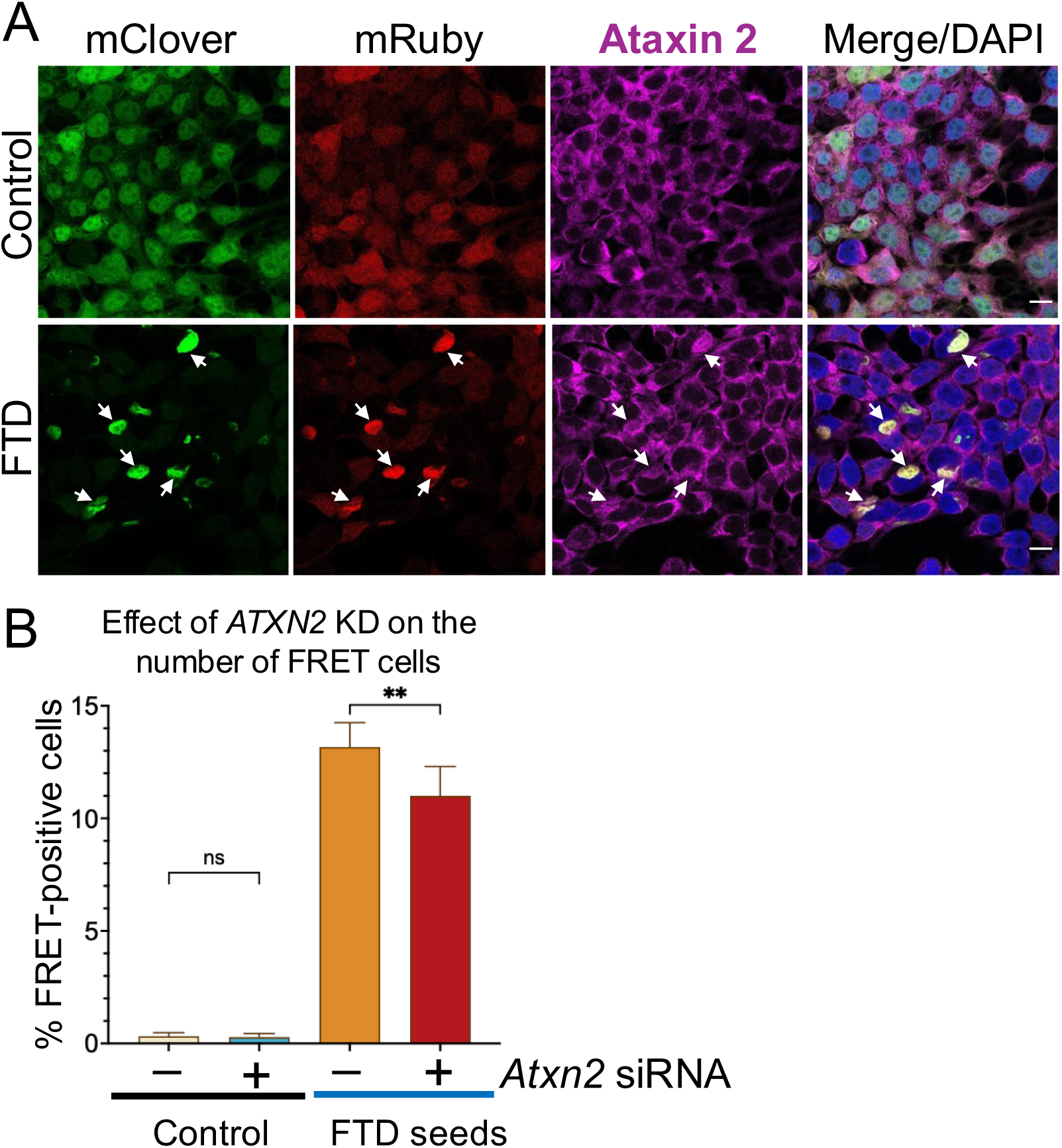
Ataxin 2 specifically colocalizes with TDP-43 aggregates triggered by seeding and promotes their accumulation. A) Representative immunofluorescence images, obtained by confocal microscopy, of HEK293^FRET^ cells treated with FTD seeds or control extract to detect ataxin 2 colocalization with TDP-43 aggregates (arrows). Scale bar, 10 μm, n>3. DAPI staining highlights nuclei in the overlay image. B) HEK293^FRET^ cells treated with FTD seeds or control extract transfected with non-targeting control or *Atxn2* siRNA after 48 hours. Cells were collected six days post-seeding, and the ratio of FRET-positive to total number of cells was quantified by flow cytometry. Values show the mean average and SEM from n>3, **p=0.009, ns, Mann Whitney test.

## DISCUSSION

TDP-43 aggregate accumulation combined with mislocalization and loss of function are hallmark features of TDP-43 proteinopathies. However, evidence linking these processes remains insufficient and the mechanisms mediating these connections are ill-defined. Our findings support a model linking these processes, whereby cytoplasmic aggregation gradually sequesters cellular TDP-43, reducing its nuclear localization and function (**Fig. 8**). We demonstrate that TDP-43 prion-like seeding is sufficient to impair autoregulation (**Fig. 5**), consistent with a decrease in active TDP-43 levels (**Figs. 3**, **4**). Our data suggest that this aberrant process establishes a toxic feed-forward loop, which may lead to uncontrolled protein expression and contribute to cytoplasmic TDP-43 aggregation. The importance of TDP-43 autoregulation in maintaining physiological proteostasis has been postulated since the discovery of this negative feedback loop^5, 53^. Its critical role in disease is further supported by data obtained from ALS/FTD patient-derived cells that lack nuclear TDP-43 and exhibit dysregulated *TARDBP* transcript expression^8^. Notably, the *TARDBP* mRNA expression profile in these cells resembles that observed in our TDP-43 aggregation model. Our findings underscore a critical role of TDP-43 autoregulation in linking aggregate accumulation with the loss of functional TDP-43, highlighting its potential as a key target in disease pathology.

**Figure 8.**
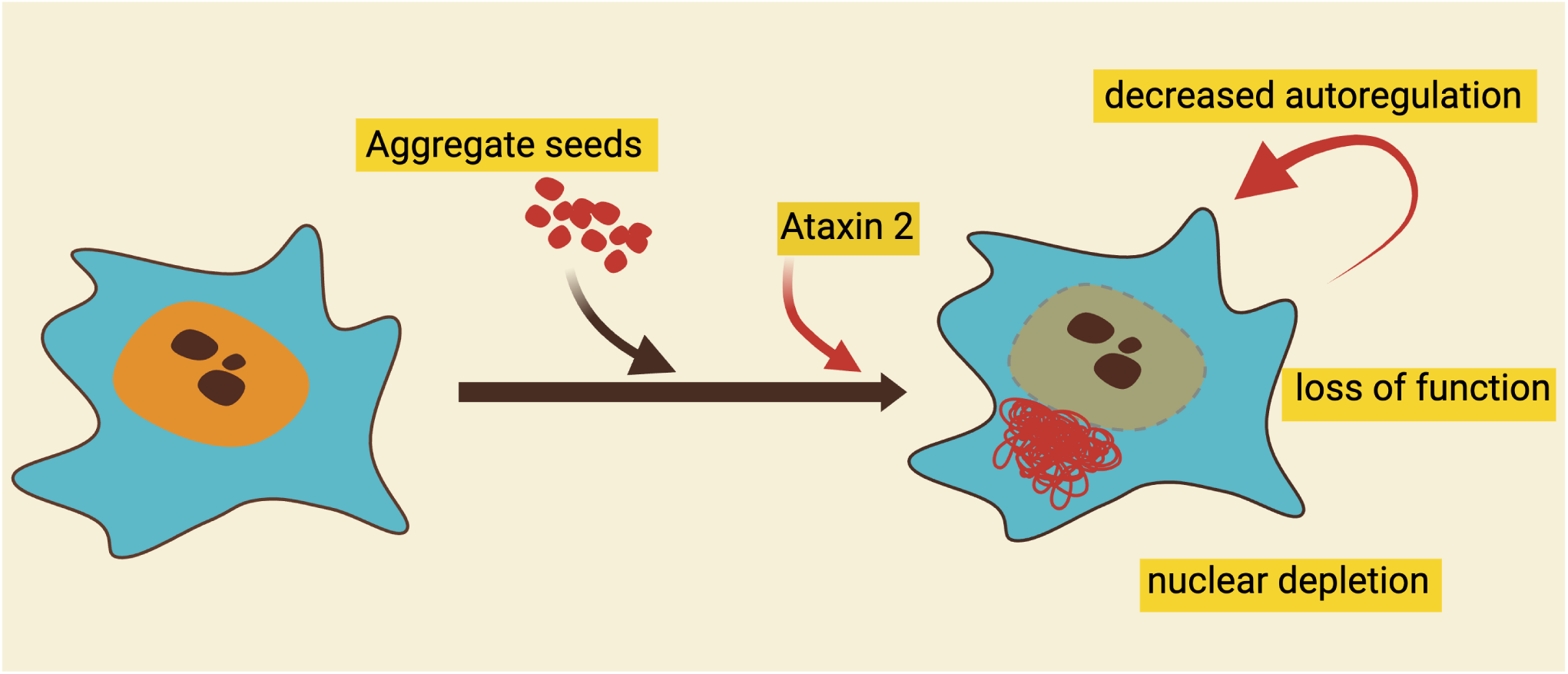
TDP-43 aggregate seeding and loss of function are linked via a toxic positive feed forward loop. Prion-like misfolding triggered by the uptake of aggregate seeds sequesters functional TDP-43, initiating a cascade of toxic events that results in progressive nuclear TDP-43 depletion, loss of function, and dysregulation of TDP-43-regulated targets. This cascade includes impaired TDP-43 autoregulation, which we propose contributes to the accumulation of misfolded protein through dysregulated protein synthesis.

Dysregulation of TDP-43 target expression in ALS/FTD patients strongly supports the idea that loss of TDP-43 function significantly contributes to neurodegeneration. Several TDP-43 regulated genes are connected to essential neuronal function and their aberrant expression may underlie disease mechanisms (e.g., *STMN2, UNC13A, KCNQ2*)^9, 10, 11, 48, 49^. In some cases, these mis-splicing events may serve as sensitive markers for TDP-43 dysfunction and disease (e.g., *HDGFL2*)^12, 50^. We surveyed the effect of aggregate seeding in the expression of mRNA transcripts regulated by TDP-43 through 3’UTR processing/alternative polyadenylation and cryptic exon splicing. Changes in cryptic exon inclusion are highly sensitive compared to alternative polyadenylation, as CE activation is detected at earlier timepoints after seeding, even when cytoplasmic CTD aggregates are observed in only 4% of cells (**Fig. 4A**). This great difference indicates that some RNA processing events regulated by TDP-43 may be more sensitive to changes in TDP-43 expression. Our findings that aggregate seeding impacts TDP-43 function are also observed in human neurons (**Fig. 4C**), indicating that this process is not cell-type specific and may be observed in disease-relevant cell types. Overall, our findings support the notion that toxic gain of function caused by TDP-43 misfolding and the loss of TDP-43 function are tightly connected processes. Importantly, this evidence strongly suggests that inhibiting TDP-43 misfolding and aggregate accumulation could mitigate TDP-43 loss of function and neurotoxicity.

By comparing the seeding properties of brain derived proteopathic seeds and aggregates generated from the full-length purified protein, we made the intriguing observation that de novo aggregates induced distinct structures suggesting that unique characteristics of nucleating seeds strongly impact the properties of the resulting aggregates. Possibly additional factors present in the brain-derived extract may also influence the structure of de novo aggregates.

These are outstanding questions relevant to elucidating mechanisms regulating TDP-43 aggregate biogenesis and toxicity, which should be addressed in future studies.

## ACKNOWLEDGEMENTS

This work was supported in part by National Institutes of Health (NIH) National Institute for Neurological Disorders and Stroke (NINDS) and National Institute on Aging (NIA) grants R01 NS114289 (Y.M.A.); and the Department of Defense CDMRP/ALSRP W81XWH-20-1-0241 (Y.M.A.). We thank the Neuropathology Core of the Knight Alzheimer Disease Research Center at Washington University in St. Louis for providing tissue used in this project.

## MATERIALS AND METHODS

All chemicals and reagents were obtained from Millipore Sigma unless otherwise specified.

### Cell culture and aggregate seeding assay

Material for aggregate seeding was obtained following sequential extraction from frontal cortex brain matter, according to previously established protocols for sequential fractionation to obtain a sarkosyl insoluble pellet^38^. Brain specimens were obtained from the Knight Alzheimer’s Disease Research Center Pathology Core at Washington University in St. Louis. Briefly, approximately 100 mg of tissue was homogenized using a mini homogenizer (Pro-PK-01200S-ProScientific) in 0.5 mL of 1% Trixon X-100 (v:v) in high salt buffer (HS, 10 mM Tris–HCl, pH 7.4, 0.5 M NaCl, 2 mM EDTA, 10% sucrose (w:v), 1 mM DTT, and protease/phosphatase inhibitor cocktail). As in all fractionation steps, the first pellet was obtained upon ultracentrifugation, 180,000*g* for 30 min at 4 °C. The first pellet was resuspended in 0.5 mL of 1% Trixon X-100 (v:v), 20% sucrose (w:v), 10 mM Tris–HCl, pH 7.4, 0.5 M NaCl, 2 mM EDTA, 1 mM DTT, protease/phosphatase inhibitors. The second pellet was solubilized in 0.1 mL of 50 mM Tris–HCl, pH 8.0, 20 mM NaCl, 2 mM MgCl2, and Benzonase (500 U/g tissue) and incubated for 20 min on ice. The third pellet was extracted with 2% sarkosyl (w:v), HS buffer and washed twice in 0.3 mL twice with phosphate buffered saline (PBS) and sonicated (Bioruptor Pico B0160010-Diagenode). The final pellet was resuspended in PBS to obtain approximately 0.1 μL/ mg of initial tissue. The amount of TDP-43 in the extracts was estimated by immunoblotting using recombinant TDP-43 as control. HEK293^FRET^ cells were generated as previously described^29^, maintained in DMEM (Dulbecco’s Modified Eagle’s Medium – High Glucose, Corning) supplemented with 10% FBS (Fetal Bovine Serum) and incubated in a humid atmosphere at 37 °C and 5% CO_2_. For the seeding assays, cells were plated in a 6-well plate at a density of 1 × 10⁵ cells per well to achieve approximately 50% confluence after 24 h. Cells were then transfected with 0.5 μL of brain-derived sarkosyl-insoluble fraction and 5 μL of Lipofectamine 2000 transfection reagent (11668-019, Invitrogen) per well in OPTI-MEM medium. After 24 hours, cells were trypsinized and replated for the indicated incubation time points.

### iNeuron differentiation, culture and seeding assays

Isogenic iPSCs derived from the KOLF2.1J parental line, obtained from the iPSC Neurodegenerative Disease Initiative (iNDI), with a stably integrated tetracycline-inducible promoter, were cultured for 3 days in the presence of doxycycline (2μg/ml). Cells were seeded at a density of 2×10^5^ cells/well in six-well plates coated with Matrigel in Dulbecco’s modified Egle’s medium (DMEM/F12) containing N2 supplement, Non-essential amino acids (NEAA), Glutamax and Y-27632. Medium was changed daily, and Y-27632 was removed after day 1. At day 4, pre-differentiated iNs were dissociated using Accutase, counted and plated at 1.5X10^6^ cells/well in 6-well plates coated with Poly-L-ornithine (0.1mg/ml) and Lamin (10ug/ml) in maturation media. Maturation media consisted of 50% DMEM/F12, 50% Brainphys neural basal media, N21Max, GDNF (10ng/ml), BDNF (10ng/ml), NT-3 (10ng/ml), Lamin (1ug/ml), doxycycline (2ug/ml). Half of the medium was replaced on day 7 and media changes were performed every 3 days thereafter using BrainPhys as the base medium. iNs were treated in 6-well plates (n=6 per condition) on day 7 with 1μl of brain-derived sarkosyl-insoluble fraction, either control or FTD seeds per well. Half of the media was replaced 3 days post-treatment, and neurons were maintained in culture for 2-3 weeks.

### Immunofluorescence microscopy

Proteins detection, visualization of mClover3 and mRuby3 signal and indirect immunofluorescence were performed according to previous methods ^44^ using the antibodies in Supplementary Table 2. Microscopy was carried out using a Keyence fluorescence microscope BZ-X series and confocal images were obtained using a Leica TCS SP8 microscope. High resolution images were obtained by stimulated emission depletion (STED) microscopy in a Leica SP8 TCS STED 3X Point Scanning Confocal microscope.

### Cell sorting and FRET quantification

Cells were rinsed and resuspended in phenol red-free DMEM for FRET quantification and FACS. FRET-positive cells were quantified using a FACSSymphony A3 flow cytometer (BD Biosciences), equipped with 405 nm, 488 nm and 633 nm lasers. The gating strategy for FRET quantification and sorting was based on analyses of stable HEK293 cells expressing mClover3-CTD or mRuby3-CTD only. This allowed us to establish the detection parameters. To measure mClover3 and FRET, cells were excited with the 488 nm laser and fluorescence was detected with a 530/30 nm and 610/20 bandpass filters, respectively. mRuby3 was excited the 561 nm laser, while emission was taken the 610/20 bandpass filter. FRET-positive cells were detected with a 610/20 bandpass filter. For each sample, we evaluated at least 30,000 cells. Data analysis was performed using FlowJo software, which quantified the percentage of FRET-positive cells based on the FRET signal intensity. FACS was performed on a FACSSymphonyA3 (BD Biosciences). Sorted cells were collected into 15 mL tubes containing DMEM; High Glucose, Corning, supplemented with 10% fetal bovine serum (FBS).

### RNA transcript expression analysis

RNA extraction was carried out using Invitrogen Purelink RNA mini kit (12183018A). RNA was treated with DNase I (EN0521-Thermo Scientific) to remove genomic DNA. RNA was extracted from iNs using the RNAeasy Mini Kit (7410-Qiagen), and genomic DNA was removed with RNase-Free DNAse set (79254-Qiagen), following the manufacturer’s instructions. cDNA synthesis and quantitative PCR were performed as previously described^67^ using primers listed in Supplementary Table 1. Glyceraldehyde-3-phosphate dehydrogenase (GAPDH) was used as housekeeping gene to normalize expression levels across samples.

### siRNA-mediated knockdown

*ATXN2* was downregulated in HEK293^FRET^ cells 48 h post-treatment with control or FTD seeds using ON-TARGETplus Human *ATXN2* siRNA-SMARTpool (Horizon Discovery L-011772-00-0005), and Non-Targeting control siRNA #1. siRNA was transfected using RNAimax (Invitrogen) according to manufacturer instructions.

---

**Supplementary Table 1:**
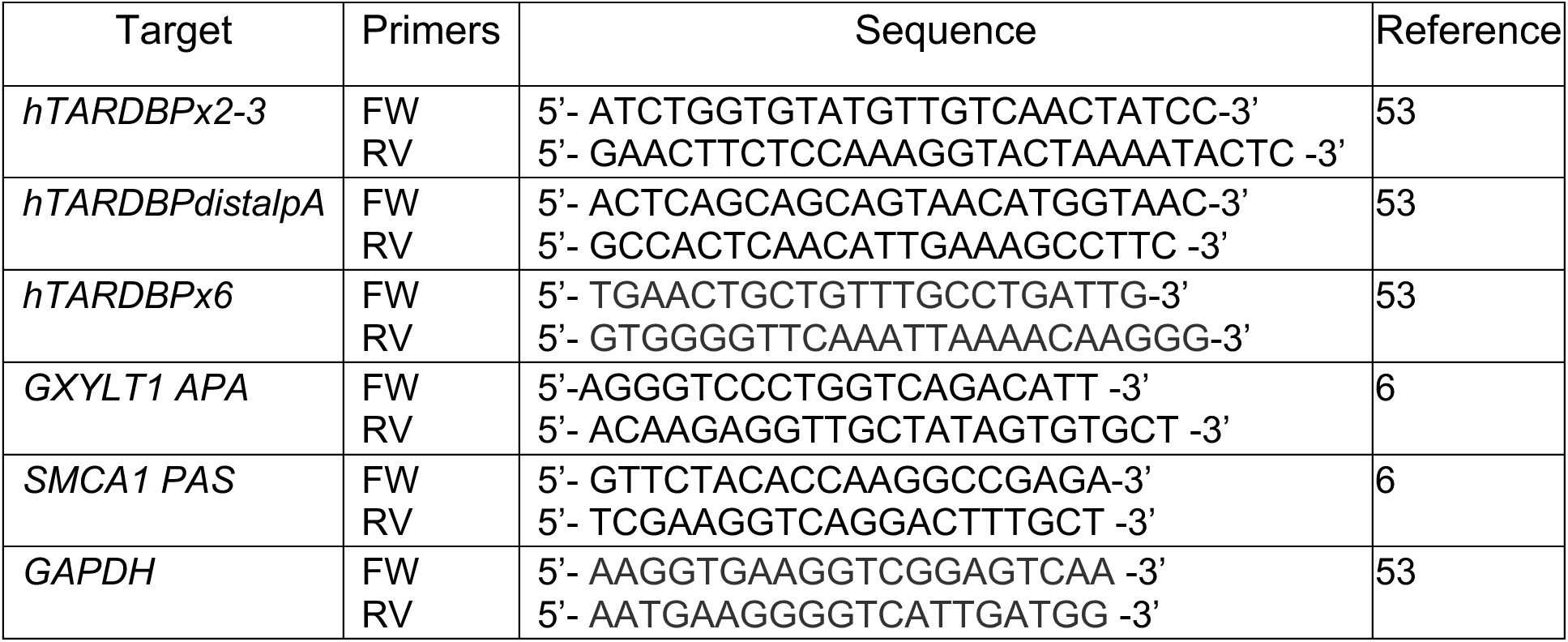
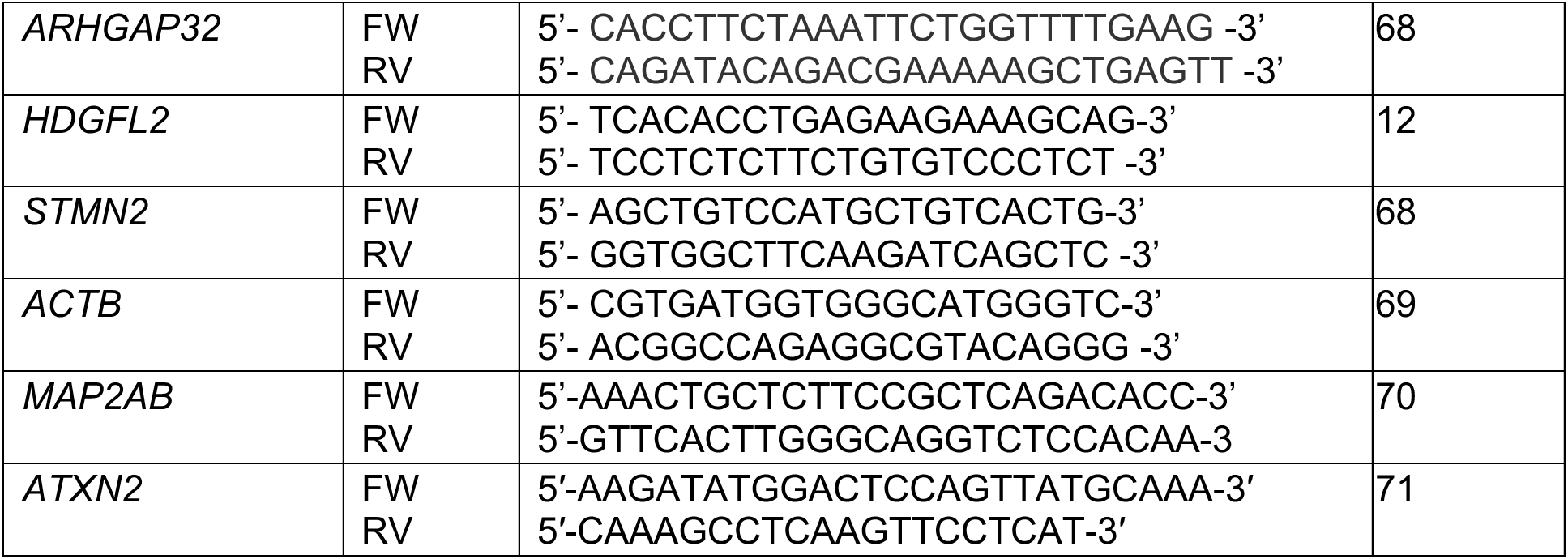
qPCR primers sequences.

**Supplementary Table 2:** List of antibodies used for immunoblotting and microscopy antibodies.

| Antibody | Reactivity | Dilution | Company | Catalog Number |
| --- | --- | --- | --- | --- |
| TDP-43 | Rabbit | IB 1:5000 | Proteintech | 10782-2-AP |
| TDP43 (C-terminal) | Rabbit | IB 1:5000 | Proteintech | 12892-1-AP |
| pTDP-43 (S409/410) | Rabbit | IF 1:1000 | Proteintech | 22309-1-AP |
| pTDP-43 (S409/410) | Mouse | IB 1:10000 | CosmoBio | TIP-PTD-M01 |
| TDP-43 (N-terminal) | Mouse | IF 1:50<br>IB 1:1000 | R&D Systems | MAB7778 |
| gH2AX | Rabbit | IF 1:200 | Cell signaling | 25775 |
| hnRNP A2B1 | Rabbit | IF 1:1000 | Abcam | Ab259894 |
| hnRNP C1/C2 | Rabbit | IF 1:250 | Abcam | Ab133607 |
| hnRNP H | Rabbit | IF 1:250 | Abcam | Ab154894 |
| Matrin3 | Rabbit | IF 1:100 | Abcam | Ab151739 |
| Ataxin 2 | Rabbit | IF 1:900 | Proteintech | 21716-1-AP |
| SFPQ | Rabbit | IF 1:50 | Abcam | Ab177149 |
| FUS | Mouse | IF 1:20 | Santa Cruz Biotechnology | Sc-47711 |
| G3BP1 | Rabbit | IF 1:200 | Proteintech | 13057-2-AP |
| p62 | Rabbit | IF 1:500 | Enzo Life Sciences | BML-PW9860 |
| GAPDH | Mouse | IB 1:10000 | Proteintech | 60004-1-Ig |

## Supplemental Figures

**Supplemental figure 1:**
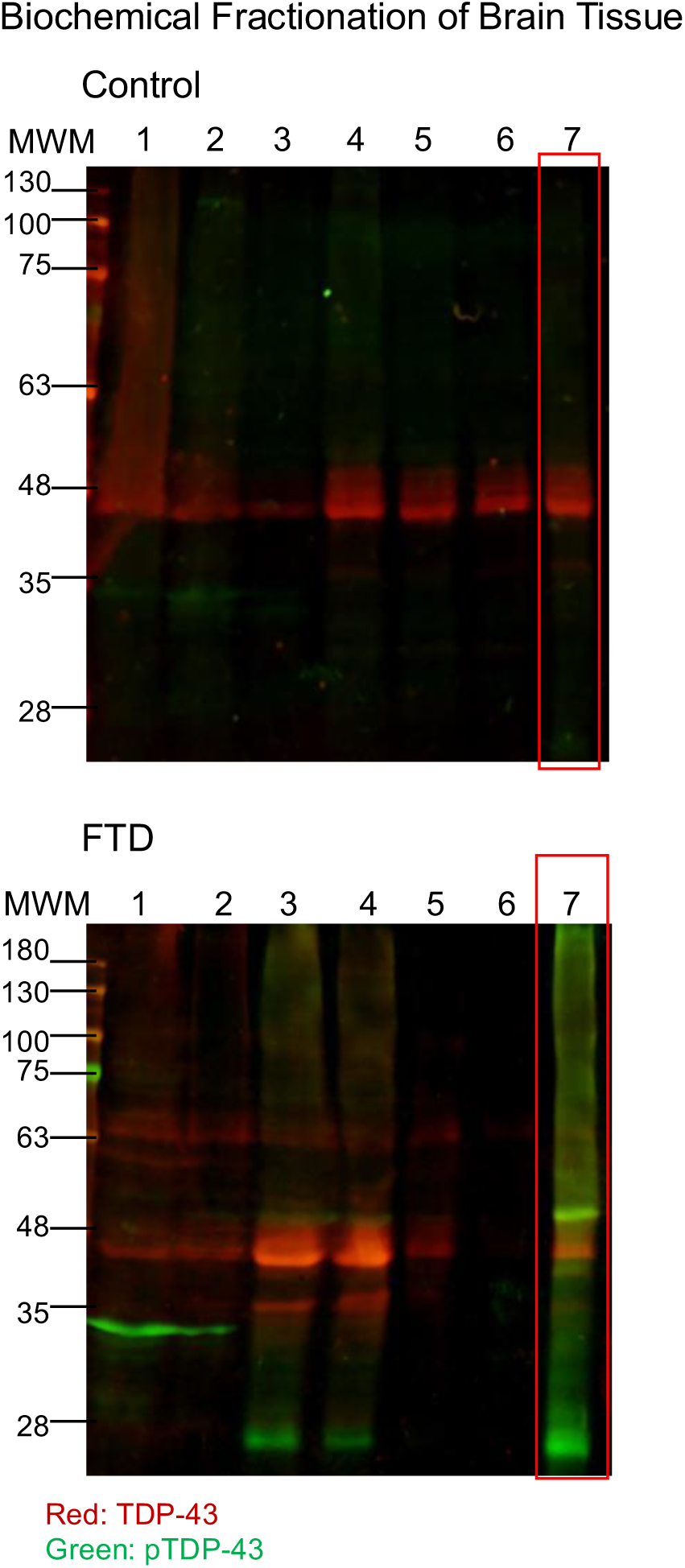
Preparation of aggregate extracts for seeding assays. Representative immunoblots of samples following fractionation^25, 38^ of brain tissue derived from control and FTD frontal cortex. Lanes were loaded with samples from each fractionation step: 1) Total tissue homogenate in high salt and Triton X-100 (HS-TX); 2) HS-TX and sucrose supernatant; 3) Supernatant of HS-TX after nuclease treatment; 4) Sarkosyl soluble samples; 5, 6) Phosphate buffered saline washes; 7) Sarkosyl-insoluble pellet.

**Supplemental figure 2:**
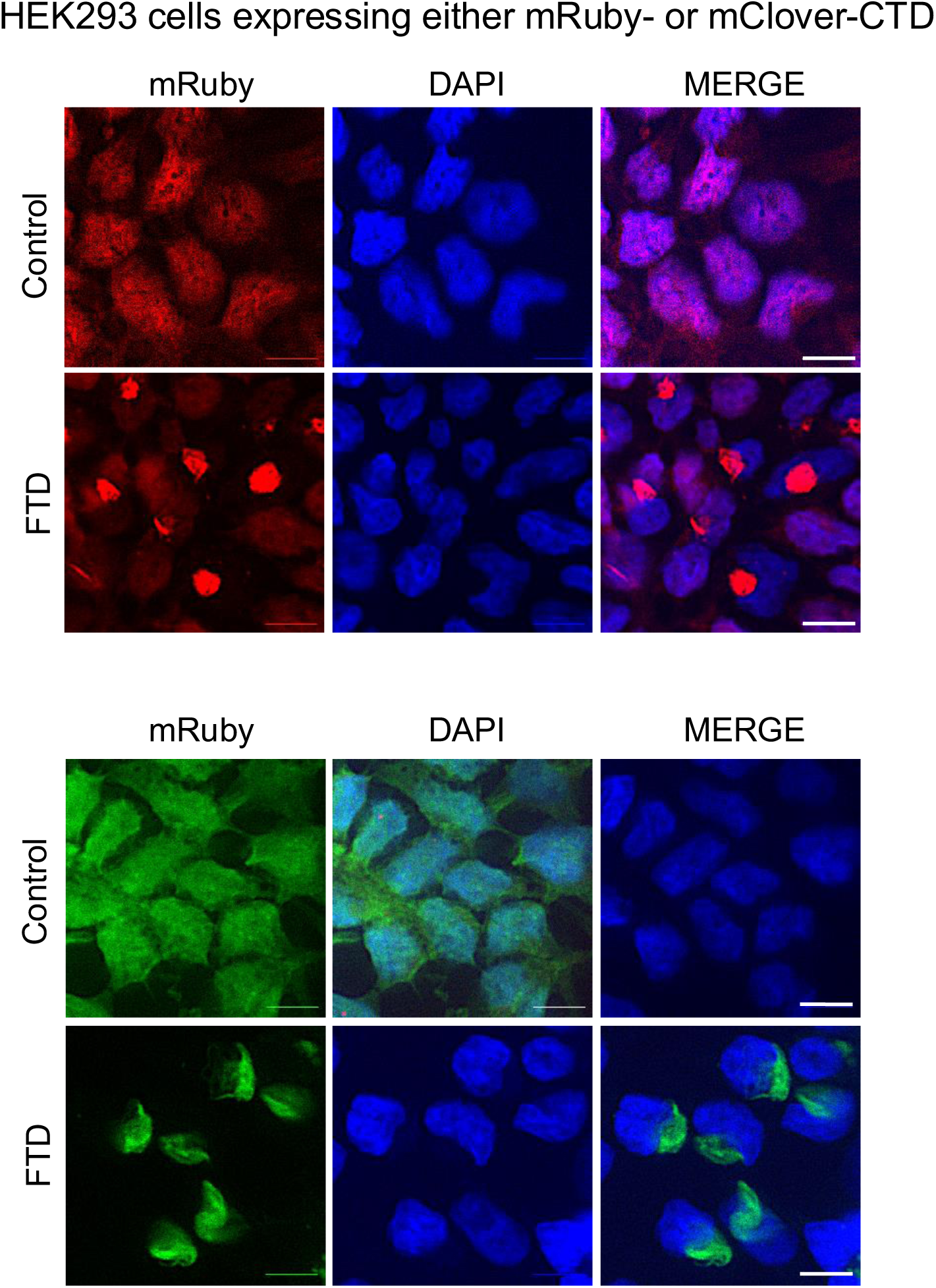
Aggregate seeding in HEK293 cells expressing only mClover or mRuby-CTD. Representative images of stable HEK293 cells expressing TDP-43 CTD N-terminally fused to mClover or mRuby treated with FTD seeds or control extract. DAPI was used to highlight nuclei. Scale, 10 μm, n>3.

**Supplemental figure 3:**
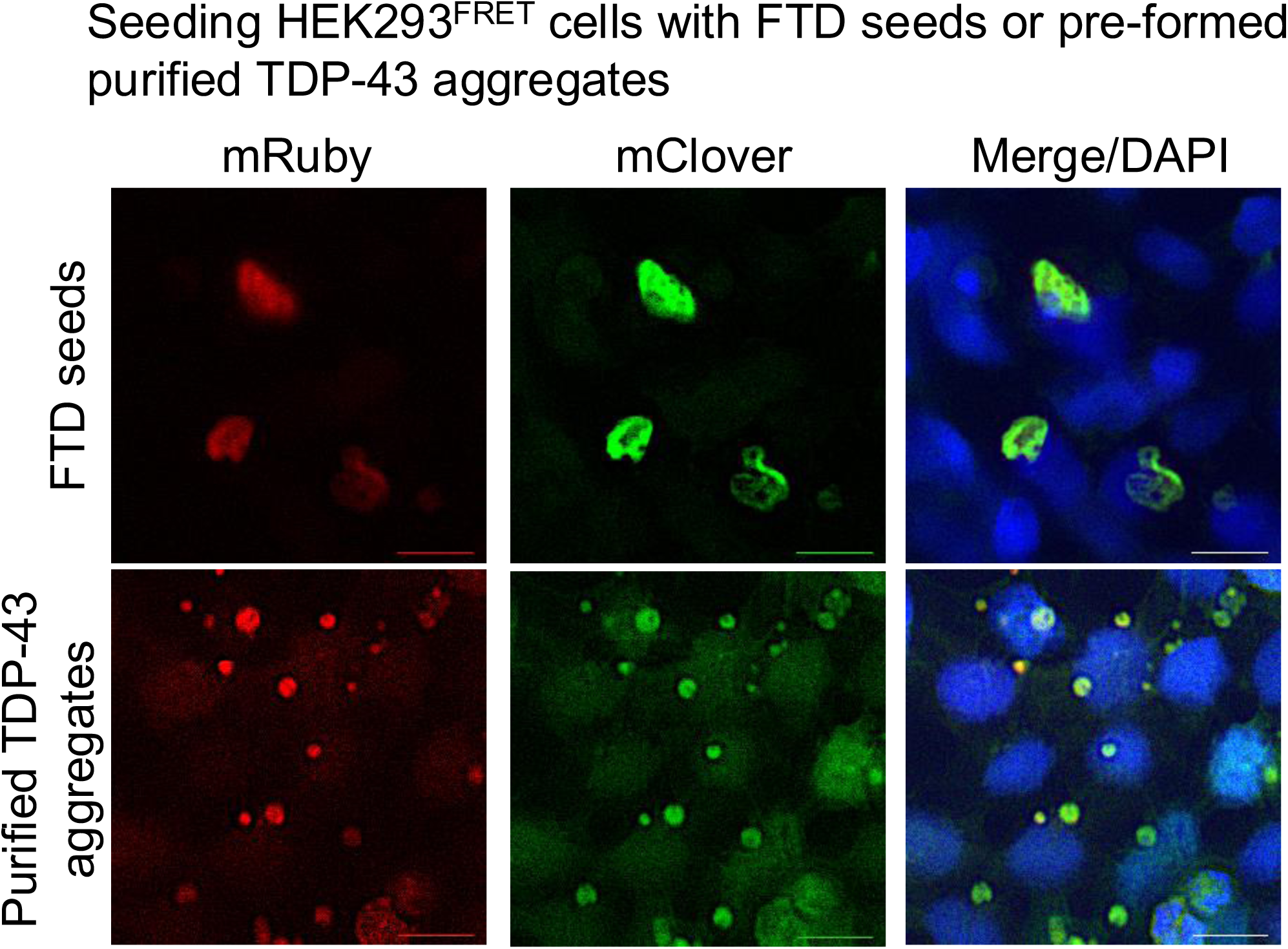
Seeds derived from patient extracts and aggregates from purified protein induce de novo aggregates with distinct morphologies. Representative images of HEK293^FRET^ cells treated with FTD seeds or pre-formed aggregates of purified full-length TDP-43^23^, imaged six days post-treatment. DAPI was used to highlight nuclei. Scale, 20 μm, n>3.

**Supplemental figure 4:**
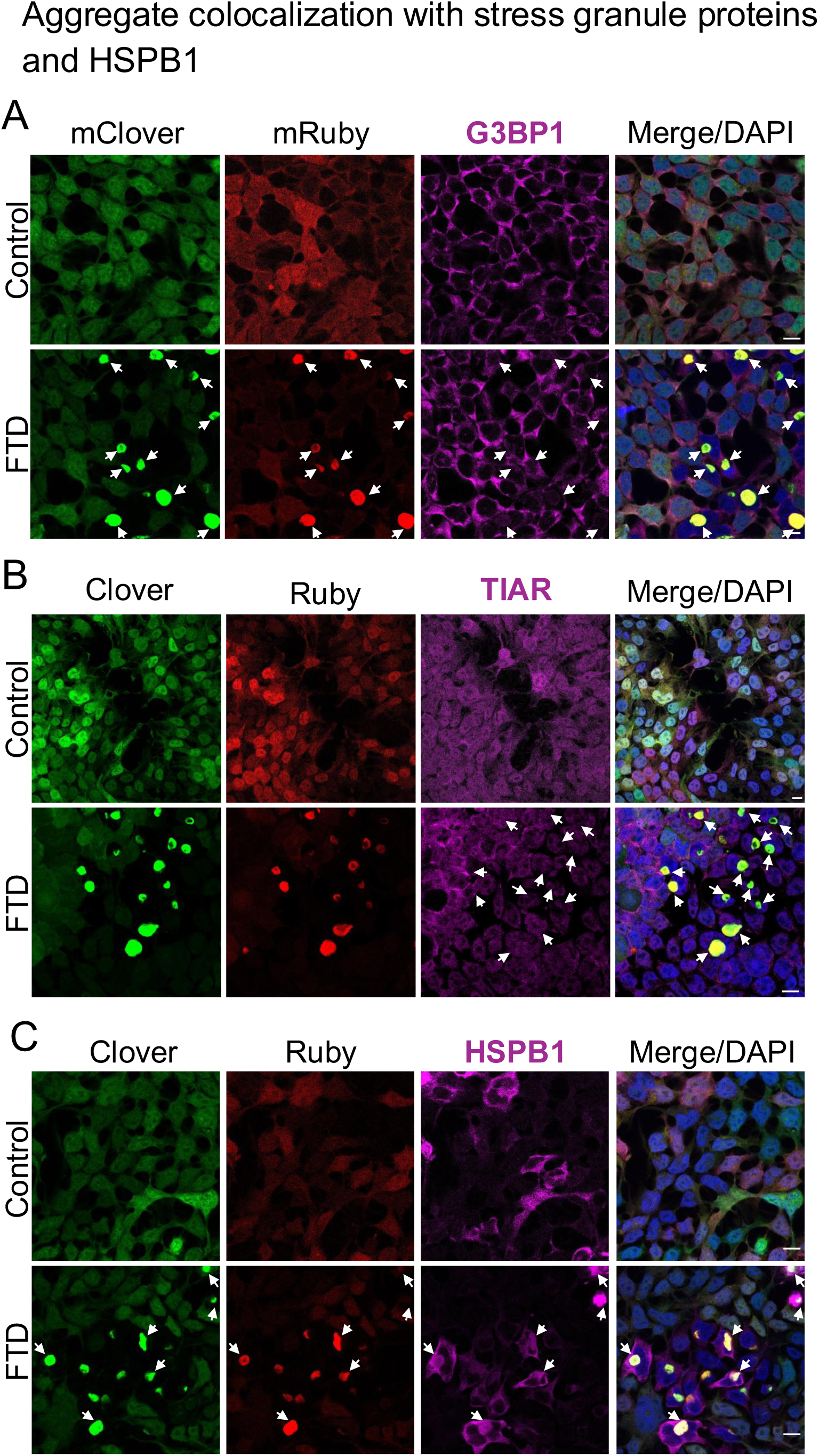
G3BP1 and TIAR, stress granule proteins, do not significantly colocalize with seeding-induced TDP-43 aggregates. Representative immunofluorescence of HEK293^FRET^ cells treated with FTD seeds or control extract, imaged six days post-treatment, using confocal microscopy. The antibodies probed for RNA binding proteins associated with stress granules G3BP1 in (A) and TIAR in (B). Arrows point to cytoplasmic TDP-43 aggregation. C) Colocalization of the small chaperone HSPB1 with TDP-43 aggregates is highlighted (arrows). Scale bar, 10 μm, n>3. DAPI staining highlights nuclei in the overlay image.

**Supplemental figure 5:**
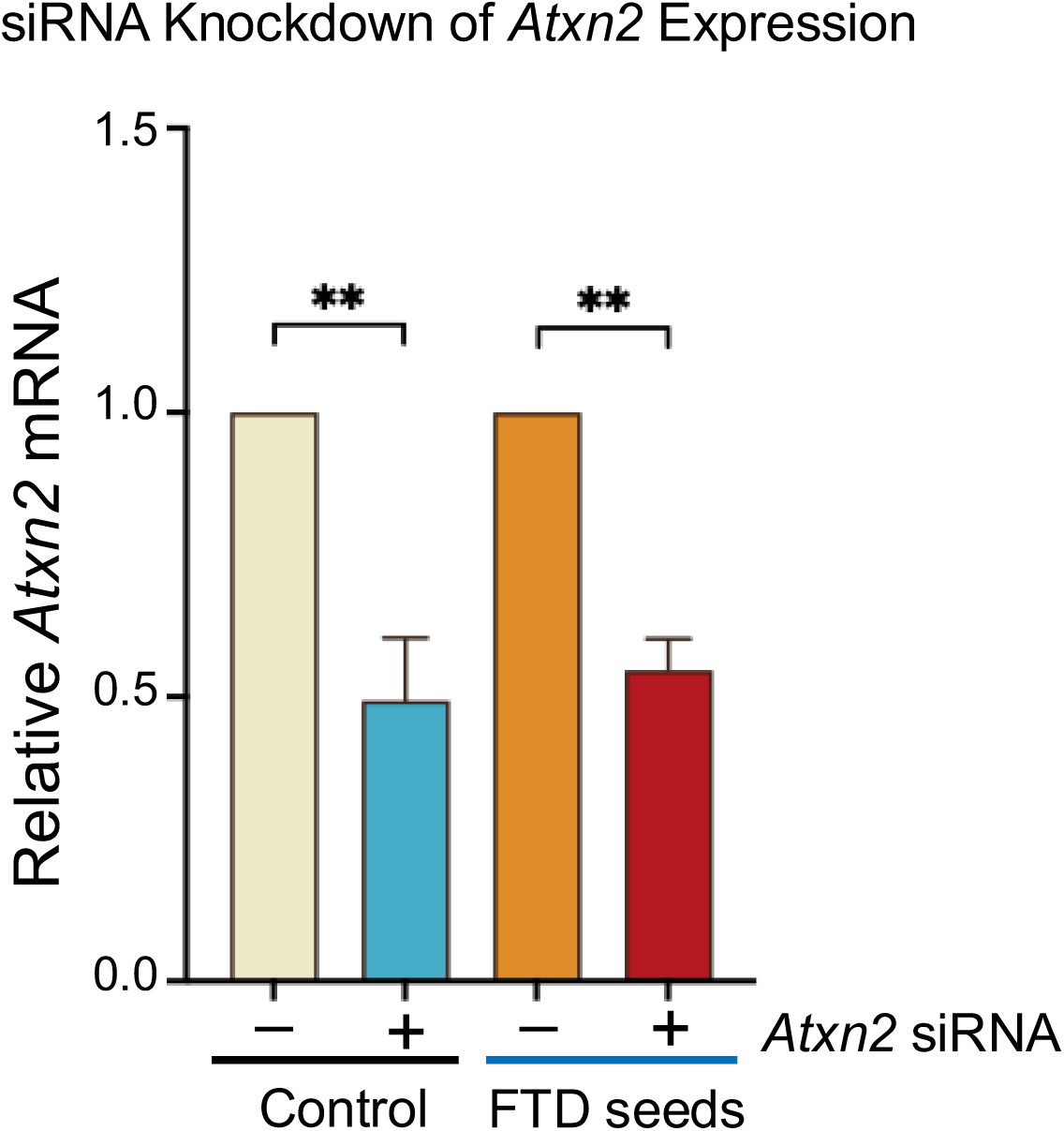
Knockdown of *Atxn2* expression. Downregulation of Atxn2 expression measured by qPCR upon siRNA treatment, compared to treatment with non-targeting siRNA control. Values show the mean average of relative mRNA expression and SEM from n>3, **p=0.009, Mann Whitney test.

## REFERENCES

1. Neumann M, et al. Ubiquitinated TDP-43 in frontotemporal lobar degeneration and amyotrophic lateral sclerosis. Science 314, 130–133 (2006).

2. Buratti E, Dork T, Zuccato E, Pagani F, Romano M, Baralle FE. Nuclear factor TDP-43 and SR proteins promote in vitro and in vivo CFTR exon 9 skipping. The EMBO journal 20, 1774–1784 (2001).

3. Mercado PA, Ayala YM, Romano M, Buratti E, Baralle FE. Depletion of TDP 43 overrides the need for exonic and intronic splicing enhancers in the human apoA-II gene. Nucleic Acids Res 33, 6000–6010 (2005).

4. Tollervey JR, et al. Characterizing the RNA targets and position-dependent splicing regulation by TDP-43. Nat Neurosci 14, 452–458 (2011).

5. Polymenidou M, et al. Long pre-mRNA depletion and RNA missplicing contribute to neuronal vulnerability from loss of TDP-43. Nat Neurosci 14, 459–468 (2011).

6. Rot G, et al. High-Resolution RNA Maps Suggest Common Principles of Splicing and Polyadenylation Regulation by TDP-43. Cell Rep 19, 1056–1067 (2017).

7. Ling JP, Pletnikova O, Troncoso JC, Wong PC. TDP-43 repression of nonconserved cryptic exons is compromised in ALS-FTD. Science 349, 650–655 (2015).

8. Liu EY, Russ J, Cali CP, Phan JM, Amlie-Wolf A, Lee EB. Loss of Nuclear TDP-43 Is Associated with Decondensation of LINE Retrotransposons. Cell Rep 27, 1409–1421 e1406 (2019).

9. Melamed Z, et al. Premature polyadenylation-mediated loss of stathmin-2 is a hallmark of TDP-43-dependent neurodegeneration. Nat Neurosci 22, 180–190 (2019).

10. Brown AL, et al. TDP-43 loss and ALS-risk SNPs drive mis-splicing and depletion of UNC13A. Nature 603, 131–137 (2022).

11. Ma XR, et al. TDP-43 represses cryptic exon inclusion in the FTD-ALS gene UNC13A. Nature 603, 124–130 (2022).

12. Seddighi S, et al. Mis-spliced transcripts generate de novo proteins in TDP-43-related ALS/FTD. Sci Transl Med 16, eadg7162 (2024).

13. Arai T, et al. TDP-43 is a component of ubiquitin-positive tau-negative inclusions in frontotemporal lobar degeneration and amyotrophic lateral sclerosis. Biochem Biophys Res Commun 351, 602–611 (2006).

14. Nelson PT, et al. Limbic-predominant age-related TDP-43 encephalopathy (LATE): consensus working group report. Brain 142, 1503–1527 (2019).

15. Amador-Ortiz C, et al. TDP-43 immunoreactivity in hippocampal sclerosis and Alzheimer’s disease. Ann Neurol 61, 435–445 (2007).

16. Josephs KA, et al. Updated TDP-43 in Alzheimer’s disease staging scheme. Acta Neuropathol 131, 571–585 (2016).

17. Josephs KA, et al. TDP-43 is a key player in the clinical features associated with Alzheimer’s disease. Acta Neuropathol 127, 811–824 (2014).

18. Meneses A, Koga S, O’Leary J, Dickson DW, Bu G, Zhao N. TDP-43 Pathology in Alzheimer’s Disease. Mol Neurodegener 16, 84 (2021).

19. McKee AC, et al. TDP-43 proteinopathy and motor neuron disease in chronic traumatic encephalopathy. J Neuropathol Exp Neurol 69, 918–929 (2010).

20. McKee AC, et al. The spectrum of disease in chronic traumatic encephalopathy. Brain 136, 43–64 (2013).

21. Brettschneider J, et al. Stages of pTDP-43 pathology in amyotrophic lateral sclerosis. Ann Neurol 74, 20–38 (2013).

22. Jamshidi P, et al. Distribution of TDP-43 Pathology in Hippocampal Synaptic Relays Suggests Transsynaptic Propagation in Frontotemporal Lobar Degeneration. J Neuropathol Exp Neurol 79, 585–591 (2020).

23. French RL, et al. Detection of TAR DNA-binding protein 43 (TDP-43) oligomers as initial intermediate species during aggregate formation. J Biol Chem 294, 6696–6709 (2019).

24. Gasset-Rosa F, et al. Cytoplasmic TDP-43 De-mixing Independent of Stress Granules Drives Inhibition of Nuclear Import, Loss of Nuclear TDP-43, and Cell Death. Neuron 102, 339–357 e337 (2019).

25. Porta S, et al. Patient-derived frontotemporal lobar degeneration brain extracts induce formation and spreading of TDP-43 pathology in vivo. Nat Commun 9, 4220 (2018).

26. Kumar ST, et al. Seeding the aggregation of TDP-43 requires post-fibrillization proteolytic cleavage. Nat Neurosci 26, 983–996 (2023).

27. King OD, Gitler AD, Shorter J. The tip of the iceberg: RNA-binding proteins with prion-like domains in neurodegenerative disease. Brain Res 1462, 61–80 (2012).

28. Arseni D, et al. Structure of pathological TDP-43 filaments from ALS with FTLD. Nature 601, 139–143 (2022).

29. Zhu J, et al. VCP suppresses proteopathic seeding in neurons. Mol Neurodegener 17, 30 (2022).

30. Lee EB, et al. Expansion of the classification of FTLD-TDP: distinct pathology associated with rapidly progressive frontotemporal degeneration. Acta Neuropathol 134, 65–78 (2017).

31. Cairns NJ, et al. TDP-43 in familial and sporadic frontotemporal lobar degeneration with ubiquitin inclusions. Am J Pathol 171, 227–240 (2007).

32. Forman MS, et al. Novel ubiquitin neuropathology in frontotemporal dementia with valosin-containing protein gene mutations. J Neuropathol Exp Neurol 65, 571–581 (2006).

33. Geser F, et al. Clinical and pathological continuum of multisystem TDP-43 proteinopathies. Arch Neurol 66, 180–189 (2009).

34. Josephs KA, Stroh A, Dugger B, Dickson DW. Evaluation of subcortical pathology and clinical correlations in FTLD-U subtypes. Acta Neuropathol 118, 349–358 (2009).

35. Mackenzie IR, et al. Heterogeneity of ubiquitin pathology in frontotemporal lobar degeneration: classification and relation to clinical phenotype. Acta Neuropathol 112, 539–549 (2006).

36. Mackenzie IR, Neumann M. Reappraisal of TDP-43 pathology in FTLD-U subtypes. Acta Neuropathol 134, 79–96 (2017).

37. Sampathu DM, et al. Pathological heterogeneity of frontotemporal lobar degeneration with ubiquitin-positive inclusions delineated by ubiquitin immunohistochemistry and novel monoclonal antibodies. Am J Pathol 169, 1343–1352 (2006).

38. Porta S, et al. Distinct brain-derived TDP-43 strains from FTLD-TDP subtypes induce diverse morphological TDP-43 aggregates and spreading patterns in vitro and in vivo. Neuropathol Appl Neurobiol 47, 1033–1049 (2021).

39. Neumann M, et al. Phosphorylation of S409/410 of TDP-43 is a consistent feature in all sporadic and familial forms of TDP-43 proteinopathies. Acta Neuropathol 117, 137–149 (2009).

40. Hiji M, Takahashi T, Fukuba H, Yamashita H, Kohriyama T, Matsumoto M. White matter lesions in the brain with frontotemporal lobar degeneration with motor neuron disease: TDP-43-immunopositive inclusions co-localize with p62, but not ubiquitin. Acta Neuropathol 116, 183–191 (2008).

41. Lynch EM, et al. Seeding-competent TDP-43 persists in human patient and mouse muscle. Sci Transl Med 16, eadp5730 (2024).

42. Conicella AE, Zerze GH, Mittal J, Fawzi NL. ALS Mutations Disrupt Phase Separation Mediated by alpha-Helical Structure in the TDP-43 Low-Complexity C-Terminal Domain. Structure 24, 1537–1549 (2016).

43. Lim L, Wei Y, Lu Y, Song J. ALS-Causing Mutations Significantly Perturb the Self-Assembly and Interaction with Nucleic Acid of the Intrinsically Disordered Prion-Like Domain of TDP-43. PLoS Biol 14, e1002338 (2016).

44. Ayala YM, Misteli T, Baralle FE. TDP-43 regulates retinoblastoma protein phosphorylation through the repression of cyclin-dependent kinase 6 expression. Proc Natl Acad Sci U S A 105, 3785–3789 (2008).

45. Mitra J, et al. Motor neuron disease-associated loss of nuclear TDP-43 is linked to DNA double-strand break repair defects. Proc Natl Acad Sci U S A 116, 4696–4705 (2019).

46. Wood M, et al. TDP-43 dysfunction results in R-loop accumulation and DNA replication defects. J Cell Sci 133, (2020).

47. Estades Ayuso V, et al. TDP-43-regulated cryptic RNAs accumulate in Alzheimer’s disease brains. Mol Neurodegener 18, 57 (2023).

48. Klim JR, et al. ALS-implicated protein TDP-43 sustains levels of STMN2, a mediator of motor neuron growth and repair. Nat Neurosci 22, 167–179 (2019).

49. Prudencio M, et al. Truncated stathmin-2 is a marker of TDP-43 pathology in frontotemporal dementia. J Clin Invest 130, 6080–6092 (2020).

50. Irwin KE, et al. A fluid biomarker reveals loss of TDP-43 splicing repression in presymptomatic ALS-FTD. Nat Med 30, 382–393 (2024).

51. Bryce-Smith S, et al. TDP-43 loss induces extensive cryptic polyadenylation in ALS/FTD. bioRxiv, (2024).

52. Fernandopulle MS, Prestil R, Grunseich C, Wang C, Gan L, Ward ME. Transcription Factor-Mediated Differentiation of Human iPSCs into Neurons. Curr Protoc Cell Biol 79, e51 (2018).

53. Ayala YM, et al. TDP-43 regulates its mRNA levels through a negative feedback loop. EMBO J 30, 277–288 (2011).

54. Koyama A, et al. Increased cytoplasmic TARDBP mRNA in affected spinal motor neurons in ALS caused by abnormal autoregulation of TDP-43. Nucleic Acids Res 44, 5820–5836 (2016).

55. Avendano-Vazquez SE, Dhir A, Bembich S, Buratti E, Proudfoot N, Baralle FE. Autoregulation of TDP-43 mRNA levels involves interplay between transcription, splicing, and alternative polyA site selection. Genes Dev 26, 1679–1684 (2012).

56. Bembich S, et al. Predominance of spliceosomal complex formation over polyadenylation site selection in TDP-43 autoregulation. Nucleic Acids Res 42, 3362–3371 (2014).

57. Weskamp K, Barmada SJ. TDP43 and RNA instability in amyotrophic lateral sclerosis. Brain Res 1693, 67–74 (2018).

58. Buratti E, Brindisi A, Giombi M, Tisminetzky S, Ayala YM, Baralle FE. TDP-43 binds heterogeneous nuclear ribonucleoprotein A/B through its C-terminal tail: an important region for the inhibition of cystic fibrosis transmembrane conductance regulator exon 9 splicing. J Biol Chem 280, 37572–37584 (2005).

59. D’Ambrogio A, et al. Functional mapping of the interaction between TDP-43 and hnRNP A2 in vivo. Nucleic Acids Res 37, 4116–4126 (2009).

60. Ling SC, et al. ALS-associated mutations in TDP-43 increase its stability and promote TDP-43 complexes with FUS/TLS. Proc Natl Acad Sci U S A 107, 13318–13323 (2010).

61. Watanabe R, et al. Intracellular dynamics of Ataxin-2 in the human brains with normal and frontotemporal lobar degeneration with TDP-43 inclusions. Acta Neuropathol Commun 8, 176 (2020).

62. Lu S, et al. Heat-shock chaperone HSPB1 regulates cytoplasmic TDP-43 phase separation and liquid-to-gel transition. Nat Cell Biol 24, 1378–1393 (2022).

63. Elden AC, et al. Ataxin-2 intermediate-length polyglutamine expansions are associated with increased risk for ALS. Nature 466, 1069–1075 (2010).

64. Wijegunawardana D, Vishal SS, Venkatesh N, Gopal PP. Ataxin-2 polyglutamine expansions aberrantly sequester TDP-43, drive ribonucleoprotein condensate transport dysfunction and suppress local translation. bioRxiv, (2023).

65. Yan X, et al. Intra-condensate demixing of TDP-43 inside stress granules generates pathological aggregates. bioRxiv, (2024).

66. Becker LA, et al. Therapeutic reduction of ataxin-2 extends lifespan and reduces pathology in TDP-43 mice. Nature 544, 367–371 (2017).

67. Koehler LC, Grese ZR, Bastos ACS, Mamede LD, Heyduk T, Ayala YM. TDP-43 Oligomerization and Phase Separation Properties Are Necessary for Autoregulation. Frontiers in Neuroscience 16, (2022).

68. Rothstein JD, Warlick C, Coyne AN. Highly variable molecular signatures of TDP-43 loss of function are associated with nuclear pore complex injury in a population study of sporadic ALS patient iPSNs. bioRxiv, (2023).

69. Choi SJ, et al. Hypoxia antagonizes glucose deprivation on interleukin 6 expression in an Akt dependent, but HIF-1/2alpha independent manner. PLoS One 8, e58662 (2013).

70. Tcw J, et al. An Efficient Platform for Astrocyte Differentiation from Human Induced Pluripotent Stem Cells. Stem Cell Reports 9, 600–614 (2017).

71. Scoles DR, et al. A quantitative high-throughput screen identifies compounds that lower expression of the SCA2-and ALS-associated gene ATXN2. J Biol Chem 298, 102228 (2022).

